# Designing a broad-spectrum four-helix bundle targeting different strains of SARS-CoV-2 Spike Receptor Binding Domains with ACE2-like binding interface

**DOI:** 10.1101/2025.06.04.657798

**Authors:** Qinghui Nie, Hongmei Yin, Ke Chen, Chao Zhang, Weiwei Li, Jianxun Qi, Jinle Tang, John Z.H. Zhang, Jian Zhan, Yaoqi Zhou

**Affiliations:** Institute of Systems and Physical Biology, Shenzhen Bay Laboratory, Shenzhen 518055, China; University of Science and Technology of China, Hefei 230026, China; Suzhou Institute for Advanced Research, University of Science and Technology of China, Suzhou 215123, China; Faculty of Synthetic Biology, Shenzhen University of Advanced Technology, Shenzhen 518055, China; Laboratory of Pathogen Microbiology and Immunology, Institute of Microbiology, Chinese Academy of Sciences, Beijing 100101, China; State Key Laboratory of Quantitative Synthetic Biology, Shenzhen Institute of Synthetic Biology, Shenzhen Institute of advanced technology, Chinese Academy of Science, Shenzhen 518055, China; NYU-ECNU Center for Computational Chemistry and Shanghai Frontiers Science Center of AI and DL, NYU Shanghai, 567 West Yangsi Road, Shanghai 200124, China; Department of Chemistry, New York University, NY, NY 10003, USA; Collaborative Innovation Center of Extreme Optics, Shanxi University, Taiyuan 030006, China; Ribopeutic Inc, Guangzhou International Bio Island, Guangdong, 510320, China

## Abstract

One major strategy for the severe acute respiratory syndrome coronavirus 2 (SARS-CoV-2) to evade antibody drugs or preventive vaccines is high mutation rate of the spike receptor binding domain (RBD). Because variable RBDs of different SARS-CoV-2 strains must bind to the same human receptor angiotensin-converting enzyme 2 (hACE2) for viral cell entry and infection, we hypothesize that designing a protein with the same or very similar hACE2 binding interface might have a broad-spectrum effect against various SARS-CoV-2 strains. The designed protein binds specifically to the WT-RBD (with micromolar affinity) but not to RBDs from other SARS-CoV-2 strains. However, two rounds of the *E. coli* display and Magnetic Cell Sorting (MACS) selection are sufficient to yield a protein named CYN1 with nanomolar binding affinities not only to the WT-RBD but also to those of Omicron BA.1, XBB.1.16, and JN.1. Molecular dynamics simulations and free-energy hotspot analysis revealed that CYN1’s broader spectrum capability stems from its engagement of essentially all ACE2 hotspot residues critical for WT-RBD binding, unlike the designed protein. The discovery of CYN1, differing by only four mutations from the designed protein, confirms that targeting the small interface of human viral receptors—rather than the entire receptor—offers a viable strategy for developing broad-spectrum inhibitors. This approach minimizes potential off-target effects arising from receptor multifunctionality.

## Introduction

Severe acute respiratory syndrome coronavirus 2 (SARS-CoV-2) quickly became a worldwide pandemic after it was first identified in December 2019. As of December 22th 2024, this virus has infected nearly 1 billion people and caused over seven million deaths according to the World Health Organization (COVID-19 cases | WHO COVID-19 dashboard, https://data.who.int/dashboards/covid19/cases?n=c). The SARS-CoV-2 virus enters the host cell through the binding of its trimeric spike protein with Angiotensin Converting Enzyme 2 (ACE2) on the human cell surface. The receptor-binding domain (RBD) on the S1 subunit of the spike protein is responsible for binding to ACE2 ^1^. The main strategy for the virus to escape the human’s immunity (either from previous infection or from the vaccines ^2–6^) is evolved mutation on the spike RBD. Many broad-spectrum antibodies and vaccines were proposed and tested but yet to be successful ^7,8^.

While the spike RBD constantly mutates, our human receptor hACE2 does not change. Thus, it is conceivable that employing hACE2 to bind with the spike RBD might offer broad-spectrum binders of spike RBDs from different virus strains. Indeed, the engineered hACE2 ^9–11^ were proposed with a demonstrated broad-spectrum effect. However, hACE2, as a hydrophobic transmembrane protein, relied on a solubilizing label ^9–13^ to be effective. Moreover, ACE2 has many functions ^14^ and direct use of ACE2 may have unintended side effects. De novo designed proteins/peptides for blocking the binding of spike to ACE2 were also proposed ^15–17^, but they are not expected to have the broad-spectrum capability of ACE2 as they have significantly modified ACE2.

Here, we designed a four-helix bundle protein based on the short 22-residue double-helical region on ACE2 that interacts with spike’s RBD. We subsequently improved it by directed evolution with a demonstrated strong binding to several spike’s RBD variants. The mechanism of this broad-spectrum capability was elucidated by conducting molecular dynamics simulations and binding free energy analysis by combining enthalpy and entropy components derived from MM/GBSA^18–20^ and interaction entropy (IE) methods^21^ with computational Alanine scanning (AS) (ASGBIE method^22–25^) for hotspot analysis^26–29^. The result confirms that improved capture of hotspot residues of ACE2 by the four-helix bundle through directed evolution is the reason behind its improved broad-spectrum capability.

## Materials and Methods

### SPalign

SPalign ^30^ is a structural alignment program developed by us that compares two protein structures using a size-independent scoring function SP-score. Here it was employed to locate the best natural backbone structure for a manually constructed four-helix bundle (H4) based on two helices in ACE2 located at the hACE2-SARS-CoV-2 RBD interface.

### OSCAR-loop

OSCAR-loop^31^ is a program for building loops for a protein region with missing amino acid residues. OSCAR-loop was kindly provided by Dr. Shide Liang. Here the program was employed to build the loop regions for the four-helix bundle H4.

### OSCAR-Design

OSCAR-Design^32^ is a protein design program that was developed by us to design protein sequences for a given protein backbone. This program employed an energy function that was based on series expansion with coefficients optimized for protein design. Here the program was employed to design protein sequences given a backbone structure and fixed sidechain conformations of RBD-binding interfacial residues copied from hACE2.

### DbD2

DbD2 (Disulfide by Design) ^33^ is a web-based program for designing disulfide bonds in proteins, which is provided at http://cptweb.cpt.wayne.edu/DbD2/. Here the program was used to add a disulfide bond to improve the stability of a designed protein as a monomer.

### Plasmids

#### Protein expressions

The H4 sequences designed and a positive control AHB2^15^ (A reported RBD binder) were optimized for *Escherichia coli* (*E. Coli*) expression system and directly synthesized and subcloned into pET-21a (+) vector (BGI Genomics, Beijing).

#### E. coli display

error prone-PCR (EP-PCR) was used to introduce mutations into a target sequence using QuickMutation™ Random Mutagenesis Kit (D0219S, Beyotime) with about 0.8% mutation rate. The mutated DNA fragments were subcloned into *E.coli* display vector pDSG323-Null-Myc (115594, Addgene) between the *Kpn*I and *Bam*HI restriction sites for construction of the mutation library.

#### Gene knockout

In order to reduce the interference of fimbriate on *E. coli* display, we knocked out the *fimA-H* from MG1655 strain using pEcCas/pEcgRNA system^34^. pEcCas (73227, Addgene) and pEcgRNA (166581, Addgen) were purchased from Addgene. The specific sgRNA sequence was designed using CRISPOR^35^ with MG1655 genetic background and then constructed into the pEcgRNA vector, named as pEcgRNA-1. The homology arm fragments (sequence in Table S6) were directly synthesized as donor DNA and subcloned into between the EcoRV and PciI restriction sites of pUC57-1.8k vector (General Biol, China).

All amino acid sequences and DNA sequences were listed in Tables S5 and S6, respectively.

### Construction of MG1655^Δ*fimA-H*^ and Generation of *E.coli* display libraries

The modified CRISPR/Cas9 system (pEcCas/pEcgRNA) ^34^ was used to knock out the *fimA-H* gene for construction of the MG1655**^Δ*fimA-H*^** strain. The pEcCas was first transformed into the MG1655 strain. Then the MG1655-pEcCas strain was co-transformed with the pEcgRNA plasmid and donor DNA by electroporation, and recovered for 1 h in 2 mL LB at 37 °C. The clones were randomly picked for colony PCR and Sanger sequencing for verifying the knockout results. Correct colonies were subjected to the plasmid curing process to obtain empty MG1655**^Δ*fimA-H*^** strains as described in the reference^34^.

The constructed library plasmid was transformed into electrocompetent MG1655**^Δ*fimA-H*^** strain by electroporation. The size of each library was 1×10^6^ clones, as determined by plating on LB-Cm agar plates with 2% w/v glucose incubated at 30 °C.

### *E. coli* display and Magnetic Cell Sorting (MACS)

We followed the previously reported *E. coli* display protocol^36^. More specifically, the *E. coli* library was induced by 1 μg/mL anhydrotetracycline (ATc) at 37 °C for overnight and harvested by centrifugation (3000 × g, 3 min). It is then washed five times with 2 mL PBS and resuspended in 1 mL of PBS. Afterwards, 100 µL of the bacteria was mixed with 50 nM Biotinylated RBD and adjusted to the final volume of 200 µL with PBS-BSA (PBS supplemented with 0.5% w/v BSA, sterile filtered and degassed), which was incubated at room temperature for 30 min. Thereafter, the bacteria were washed five times with 1 mL of PBS-BSA, resuspended in 100 µL of the same buffer containing 20 µL of anti-biotin paramagnetic beads (Miltenyi Biotec), and incubated at 4 °C for 20 min. Next, the bacteria were washed three times with 1 mL of PBS-BSA and resuspended in 500 µL of the same buffer, of which 10 µL was kept aside to calculate the input bacteria before the MACS procedure and the remaining portion (490 µL) was applied onto a MACS MS column (Miltenyi Biotec), that was previously equilibrated with 500 µL of PBS-BSA and placed on the OctoMACS Separator (Miltenyi Biotec).

The unbound cells (flow-through) were collected, and the column was washed five times with 2 mL of PBS-BSA. The wash was combined with the flow-through as the “Unbound fraction”. Next, the column was removed from the OctoMACS Separator and placed onto a new collection tube to free the bound fraction by washing with 2 mL of PBS-BSA. The bound fraction was applied onto a new equilibrated MACS MS column for reduced bacterial non-specific binding. The column was washed five times with 2 mL of PBS-BSA^36^. Next, the low-affinity mutants were washed away by 10 mM Glycine-HCl pH 2.0 and pH 3.0 (two times with 2 mL each time, and three times for second-round evolution), and the high-affinity mutants were still on the column. Afterwards, the column was removed from the OctoMACS Separator and placed onto a new collection tube, 2 mL of LB was added, and the cells were eluted out. This fraction was labeled as the “Screen fraction”. Serial dilutions of screen fractions were plated to determine CFU and perform the Sanger sequencing.

### Expression and purification of recombinant proteins

Expression plasmids were transformed into chemically competent *E. coli* Shuffle T7-B. Protein expression was then induced with 0.1 mM of isopropyl β-D-thiogalactopyranoside (IPTG) at 16 ℃. After overnight induction, the cells were harvested by centrifugation at 10,000 × g for 30 min and then resuspended in lysis buffer (150 mM NaCl, 20 mM Tris-HCl (pH 8.0), with 0.1 % triton X-100). The cells were lysed with an Ultrasonic Homogenizer (JY92-IIN, Scientz) for 20 minutes (2 sec on-3 sec off) at an amplitude of 30 %, and then centrifuged at 30,000 × g for 30 min. The precipitate was then removed, and the supernatant was collected. The supernatant was purified by affinity chromatography (HisTrap FF crude 5 mL, Cytiva) and followed by size exclusion chromatography (HiLoad 16/600 Superdex 75 pg, or Superdex 75 Increase 10/300 GL, Cytiva) on the ÄKTA pure™ 25M system (Cytiva). All protein samples were characterized using 4-20 % gradient SDS-PAGE and their purity was higher than 95%. Protein concentrations were determined using a NanoDrop spectrophotometer (Thermo Scientific) with absorbance at 280 nm using predicted extinction coefficients (https://web.expasy.org/protparam/).

### Circular dichroism spectroscopy

Far-ultraviolet CD measurements were carried out with a JASCO-1500 equipped with a temperature-controlled multi-cell holder. Wavelength scans were recorded from a wavelength region of 198 - 255 nm at 25 ℃, 95 ℃, and 25 ℃ after melt, and taking the average of four repeats. Temperature melts monitored dichroism signal at 222 nm in steps of 2 ℃/minute from 16 to 95 ℃ with 30 s of equilibration time ^15^. Wavelength scans and temperature melts were performed using 0.025 mg/mL protein in PBS buffer (1.8 mM KH_2_PO_4_, 10 mM Na_2_HPO_4_, 137 mM NaCl, 2.7 mM KCl, pH 7.4) with a 1 mm path-length cuvette.

Melting temperatures were determined by fitting the data with a sigmoid curve equation by the instrument-integrated software. Helical contents were analyzed according to the reported method^37^: negative ellipticity at 208 and 222 nm of the CD spectrum. The α-helical content was predicted using K2D3 web server (<u>K2D3:</u> <u>Estimating protein secondary structure from CD spectra</u>) ^38^ with the following parameters: (1) the CD spectra from 198 to 240 nm was converted to molar residue ellipticity (MRE, deg·cm²·dmol⁻¹) units prior to analysis; (2) structural prediction constraints include specifying protein length as 146 amino acid residues.

### Biolayer interferometry

Biolayer interferometry (BLI) binding data were collected in Octet RED384 (ForteBio) at 30 ℃ and processed using the instrument-integrated software. All RBDs were biotinylated by the biotinylating kit (GENEMORE, Cat. G-MM-IGT) according to the kit’s manual. The biotinylated RBDs were loaded onto streptavidin-coated biosensors (SA ForteBio, Cat. 18-5019) in PBS (pH 7.4) buffer. The buffer of analyte proteins was exchanged to binding buffer (PBS, 0.05% surfactant P20) by Amicon® Ultra Centrifugal Filter (Millipore) and then diluted to the desired concentrations. After baseline measurement with the binding buffer alone, the binding kinetics were monitored by dipping the biosensors in wells containing the target protein at the indicated concentration (association step) and then dipping the sensors back into baseline/buffer (dissociation). The data were processed using ForteBio Data Analysis software 12.0, and the equilibrium dissociation constant (*K*_D_) values were obtained by global fitting with 1:2 and 1:1 binding models.

### SARS-CoV 2 RBDs

The RBDs of SARS-CoV 2 variants (WT, XBB.1.16 and JN.1) were purchased from Sino Biological (Cat. 40592-V08B, 40592-V08H136, 40592-V08H155), and the RBD of BA.1 was kindly provided by Haitao Yang’s Group. We chose the Omicron BA.1, XBB.1.16 and JN.1 as the representatives of virial strains for testing our designed proteins because these strains were a part of the important SARS-CoV-2 evolution nodes, and have a high immune evasion ability to the anti-spike antibodies or vaccines that were developed based on their strain’s ancestors ^7^.

### Molecular Dynamics Simulations

The complex structure of ACE2 and WT SARS-CoV-2 Spike RBD (WT-RBD) was obtained from PDB (PDB ID: 6M0J). As each designed protein has two binding interface to WT-RBD, the complex structures of WT-RBD with D2-133 in either its helices 1-2 or helices 3-4 regions were predicted by OSCAR-Design (Figure 1). Other complex structures of CYN1, E6, and E9 in their helices 1-2 or helices 3-4 regions with WT-RBD were mutated from D2-133. All complexes were filled by the leap module in AMBER20 package^39^. All MD simulations were performed using pmemd.cuda in AMBER20 with ff14SB force field for the protein in a truncated octahedron box formed by TIP3P explicit water molecules. The closest distance between any atoms originally present in solute and the edge of the periodic box was 12 Å. The particle mesh Ewald (PME) was used to treat the long-range electrostatic interactions. The nonbonded interactions were truncated with 10 Å cutoff. Periodic boundary condition (PBC) was imposed on the system during the calculation of nonbonded interactions. The time-step was set at 2 fs and SHAKE^40^ was used to constrain the bonds involving hydrogen atom. Langevin thermostat with the collision frequency of 2.0 was applied to control the temperature. Firstly, the system was minimized with the protein constrained to equilibrate the solvent. Then, the protein was released to minimize the whole simulation system. Afterward, the system was slowly heated to 300 K, followed by a 40 ns equilibration of the whole system in an NPT ensemble at an interval of every 10 fs, and during this session, no restrain was performed. Then, a 10ns production run was performed to obtain 5000 snapshots for free-energy and hotspot analysis by ASGBIE.

**Figure 1.**
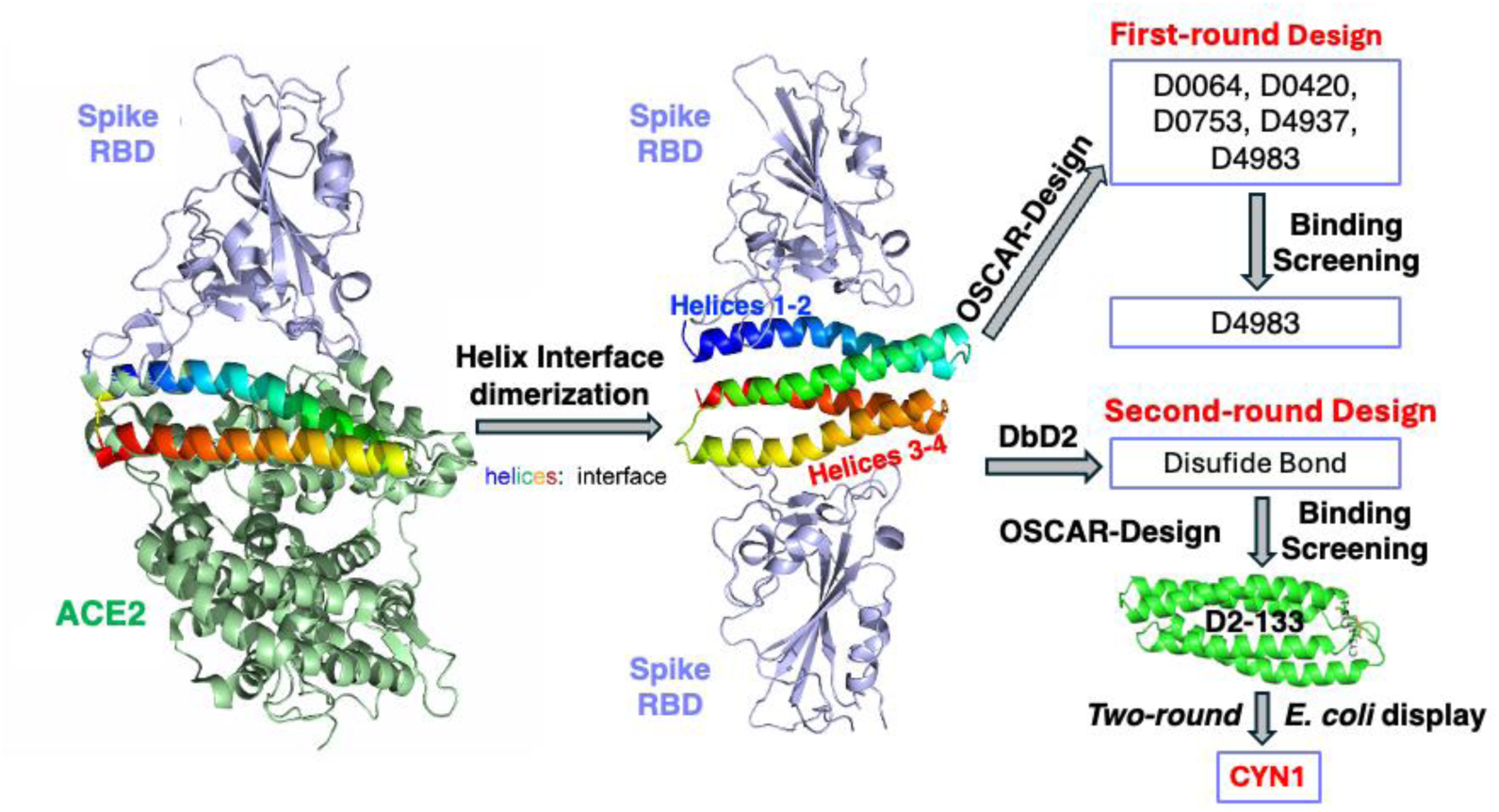
The flowchart of the iterative design and improvement of a four-helix bundle (H4). The designed protein has two copies of the two-helix binding interface of hACE2 (helices 1-2 and 3-4) to SARS-CoV-2 Spike receptor binding domain (RBD), whose backbone structure was modeled after the closest four-helix template in the protein databank. The sequences for this backbone structure along with fixed binding residues were designed by OSCAR-Design and five were tested for binding affinity. To improve monomeric stability, the second-round design was performed after adding a disulfide bond. The best design (D2-133) was improved by directed evolution using *E. coli* display and Magnetic Cell Sorting (MACS) to obtain the final protein CYN1.

### Free-energy and hotspot analysis by ASGBIE

The ASGBIE method^22–25^combines enthalpy and entropy components derived from MM/GBSA^18–20^ and interaction entropy (IE) methods^21^ with computational Alanine scanning (AS) for hotspot analysis^26–29^. Here, we simply followed the same method utilized in Ref. ^22–25^.

## Results

The flowchart of the overall design, subsequent screening and redesign as well as final improvement of four-helix ACE2-interface mimics by directed evolution is shown in Figure 3. Specific details are described as below.

### Initial design of a four-helix bundle (H4) with two ACE2-Spike binding interfaces

#### Backbone Structure

We obtained the interface residues between human ACE2 (hACE2) and wild-type SARS-CoV-2 Spike protein according to their complex structure (PDB 6M0J) based on contact distances and the reduction of solvent accessible surface area (SASA) per residue after complex formation. Most interface residues in hACE2 were found located on the two N-terminal helices. After careful inspection of the conformation of the two helices, we built a four-helix bundle prototype by manually dimerizing the two N-term helices of hACE2. This manually built dimeric structure together with binding Spike proteins was further relaxed using ROSETTA^41^. The relaxed structure was then compared to all existing four-helix bundle structures in RCSB PDB with SP-align^30^. All structurally aligned PDBs were sorted according to the SPe-alignment score and top 10 ranked structures were manually inspected and the structure of the serine-rich domain from Crk-associated substrate (PDB 1Z23) was found as the closest match and employed to adjust the backbone structure of the prototype by modifying the relative distance and orientation between dimerized two-helix bundles.

#### Sequence Design

We copied the conformations of ACE2 interface residues onto the prototype of the four-helical bundle: residue 22-84 from ACE2 in 6M0J were used to build the N-term half of the four-helix bundle, with P84 modified to G84, and residues 22-88 from ACE2 in 6M0J were extracted to build the C-term half. Residues 85-88 were included in building the C-term half, because they mimicked well the terminal cap of the template H4 in PDB 1Z23, which may contribute to the stability of designed H4. The gap between the two halves was connected by building the loops of various lengths (5-9) modeled using OSCAR-loop ^31^. Note that residue 84 from N-term and residue 22 from C-term were freely mutated during loop modeling and were included in loop modelling. All residues were then re-numbered starting from 1. The H4 structure was built to have two binding interfaces to interact with the Spike protein. The above constructed structures with different loop lengths were further optimized with AmberTools ^42^ and inspected with Procheck ^43^. One backbone structure, which has a modeled loop of 9 residues passed the check. Thus, this backbone structure was employed to redesign by OSCAR-Design ^32^ with 2x22 fixed interface residues (N-term half: E2, Q3, T6, F7, D9, K10, H13, E14, E16, D17, Y20, Q21, L24, N40, N43, K47, F51, E54, Q55, L58, M61 and Y62; C-term half: E72, Q73, T76, F77, D79, K80, H83, E84, E86, D87, Y90, Q91, L94, N110, N113, K117, F121, E124, Q125, L128, M131 and Y132) in the presence of two fixed Spike proteins (Figure 1).

#### Sequence Selection

Five thousand sequences designed by OSCAR-Design were clustered with CD-HIT ^44^ and filtered according to the area of the largest hydrophobic surface-patch 𝑆_𝐻𝑃𝐴_, the number of low-complexity areas 𝑁_𝐿𝐶𝐴_ and the number of atomic clashes 𝑁_𝑐𝑙𝑎𝑠ℎ_. The areas of hydrophobic surface-patches were calculated by QUILT ^45^. The number of low-complexity areas was calculated by segmasker from the NCBI BLAST+ package ^46^. We found that there were five sequences with sequence identity lower than 75% between each other, 𝑆_𝐻𝑃𝐴_ ≤ 550Å^2^ (509 Å^2^ for the original sequence), 𝑁_𝐿𝐶𝐴_ = 0 and 𝑁_𝑐𝑙𝑎𝑠ℎ_ = 0. These sequences (D0064, D0420, D0753, D4937, and D4983) were selected for the first-round experimental test (Table S1). Three of the five designs (D0753, D4937, and D4983) were expressed soluble and purified (Figure S1). All have micromolar binding affinities to WT-RBD with D4983 having the strongest binding affinity (Figure S2). However, these three designed proteins all existed in aggregated forms on a chromatographic column (Figure S3).

#### Adding Disulfide-bond and Redesigning Sequences

To make the monomeric folding of the designed proteins more stable, we introduced one disulfide bond to the backbone structure in its non-interfacial areas by using the program DbD2 (<u>Disulfide</u> <u>by Design</u>) ^33^. This process produced several designs with the disulfide bond added to various locations. These designs were further structurally optimized and sequence-redesigned in the same way as in the first-round design with the disulfide-bond fixed. We collected 1000 designs and further filtered with 𝑆_𝐻𝑃𝐴_ ≤ 385Å2, 𝑁_𝐿𝐶𝐴_ = 0, and 𝑁_𝑐𝑙𝑎𝑠ℎ_ = 0. This results in six second-round sequences with disulfide bonds located at CYS1-CYS133 (D2-133), CYS22-CYS48 (D2-128), CYS25-CYS44 (D2-477), CYS70-CYS136 (D2-247), CYS92-CYS118 (D2-830) and CYS95-CYS114 (D2-099), respectively (Figure 1 and Supplementary Table S2). Among 6 new designs (D2-099, D2-128, D2-133, D2-247, D2-477, D2-830), only D2-133 was monomeric, D2-099, D2-477 were partly monomeric, whereas D2-128, D2-247, and D2-830 were mostly multimeric (Figure 2A, 2B and 2C). Thus, only D2-133 was employed for the binding affinity test.

**Figure 2.**
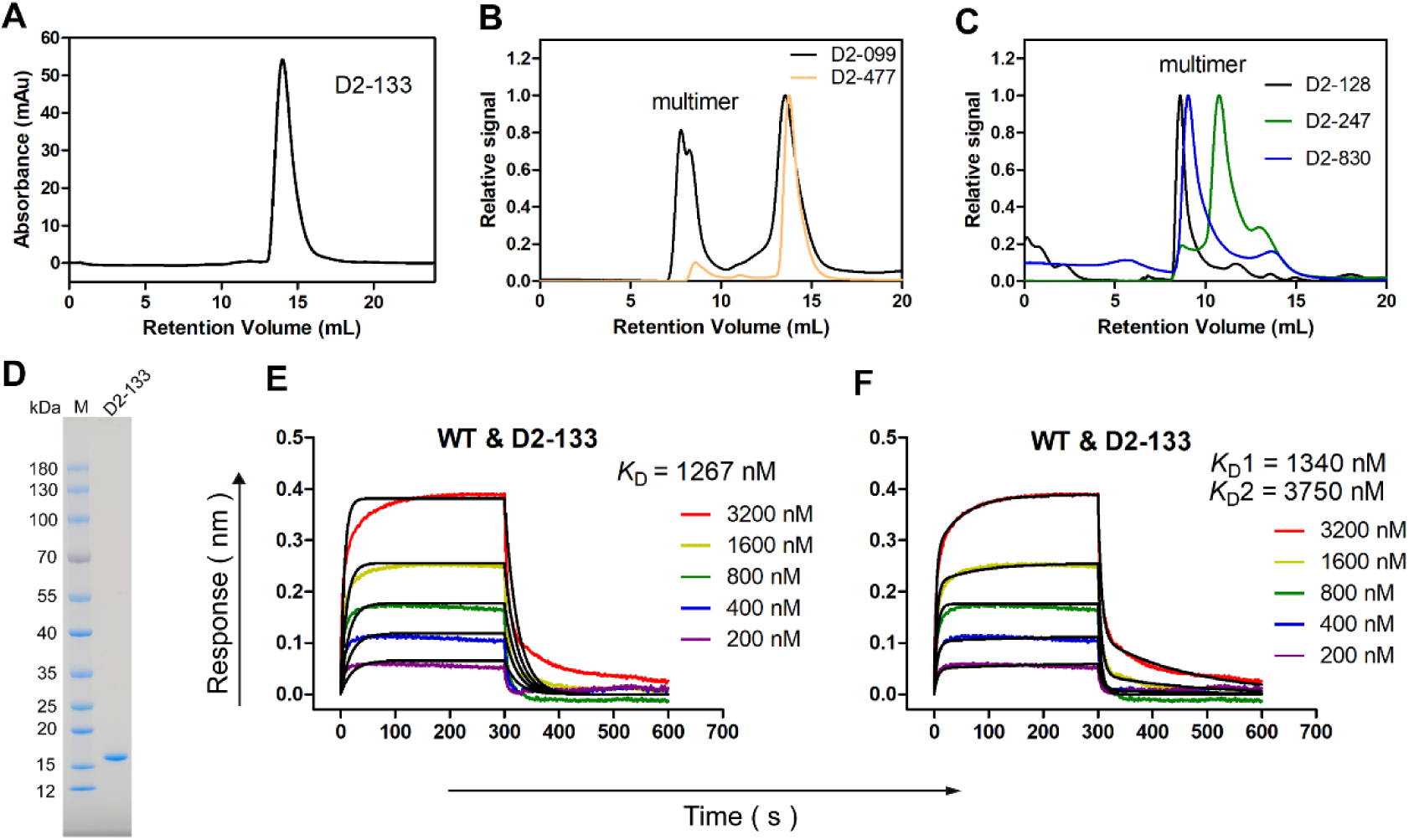
Expression and purification of the second-round designs of H4 along with affinity measurement. Size-exclusion chromatography for D2-133 (**A**), D2-099 and D2-477 (**B**), and D2-128, D2-247, and D2-830 (**C**) indicates that only the D2-133 design is purely monomeric. (**D**) Purified D2-133 was shown on a 4-20% SDS-polyacrylamide gel and stained with Coomassie brilliant blue. The binding affinity between D2-133 and WT SARS-CoV-2 Spike RBD at the micromolar level according to the Biolayer-interferometry (BLI) measurement (Fitted by 1:1 model (**E**) and 1:2 model (**F**)).

### The designed ACE2 mimic D2-133 exhibited micromolar affinity against RBD domain of Spike protein

The purified monomeric D2-133 was obtained from Size-exclusion chromatography (Figure 2D). Its binding affinity to RBD of wild type Spike was determined using BLI (Figure 2E and 2F). The D2-233 showed a micromolar affinity (1.3 μM). This affinity was similar to the AHB2 (affinities ranging from 100 nM to 2 μM) ^15^ or DBR3_02 (*K*_D_: 4.6 μM), previously designed RBD binders ^17^. Compared with the 1:1 fitting model (Figure 2E), D2-133 showed a better fitting with the 1:2 fitting model (Figure 2F), consistent with our inclusion of two hACE2-RBD binding interfaces from hACE2.

### Improving D2-133 by *E. coli* display and MACS selection

The *E. coli* display was used to improve the D2-133 affinity. To exclude the influence of fimbriae on display^36^, *fimA-H* was knocked out from MG1655 using pEcCas/pEcgRNA system ^34^ and the MG1655^Δ*fimA-H*^ strain was successfully constructed for bacterial display (see Figure S4 and Methods). EP-PCR was used to introduce mutations on the D2-133 sequence for the construction of *E. coli* display library with about 0.8% mutation rate.

The D2-133 library can bind to RBD-labeled beads but not to empty beads as shown in Figure 3A (Group 1 vs Group 2). However, neither strong acid (Figure 3A, Group 3) nor high salt (Figure 3A, Group 4) conditions can remove low-affinity mutants from the library. Thus, we increased the selection pressure by lowering the pH in round 1 (Figure 3B, Group C and D). Two clones (labeled as E-6 and E-9) from round 1 were selected for expression and purification for affinity evaluation using BLI. Both E-6 and E-9 have double mutations over D2-133 (K70E and K90R for E-6 and N103I and E125G for E-9). Both improve over D2-133 in binding affinity to WT-RBD with *K*_D_ of 99 nM and 501 nM, respectively (Figure 3B and 3C).

**Figure 3.**
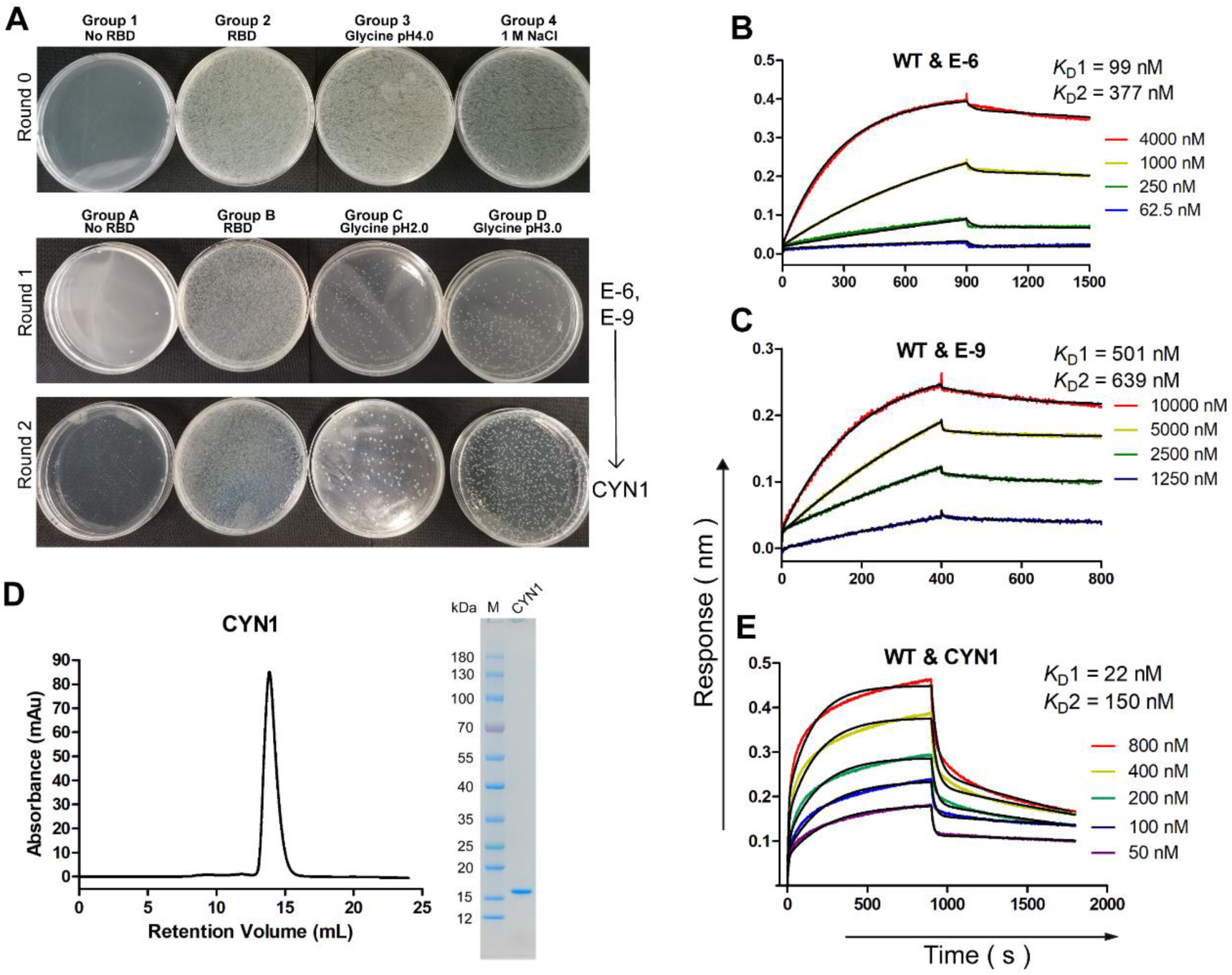
Improved binding affinity of D2-133 by *E. Coli* display and MACS. (**A**) Round 0 indicates poor selection of D2-133 mutants with pH 4.0 (Group 3) and 1M NaCl (Group 4), compared to no selection pressure (Group 2). Increased selection pressure at pH 2.0 and 3.0 in Round 1 led to E-6 and E-9. In Round 2, the selection of E-6 mutants led to CYN1. (**B**) and (**C**) The binding affinities of E-6 and E-9 with WT-RBD were measured by BLI. (**D**) Size-exclusion chromatography for CYN1. (**E**) The binding affinity of CYN1 with WT-RBD measured by BLI.

Employing the best-affinity sequence E-6, we performed the second round of *E. coli* display and MACS selection (Figure 3A). One clone CYN1 was selected. It has four mutations over D2-133 (L59T, K70E, K90R, L129T). CYN1 was expressed and purified in the monomeric state (Figure 3D). It has a binding affinity of 22 nM to WT Spike RBD, about 60 times stronger than that of D2-133 (Figure 2 F vs Figure 3 E) and 4 times stronger than that of hACE2 with WT Spike RBD ^47–49^.

### High thermal stability of CYN1

To investigate the secondary structural content and thermal stability of D2-133 and CYN1 at far-UV region, Circular dichroism (CD) experiments were carried out using Circular Dichroism Spectrophotometer J-1500. The results of CD spectrum indicated D2-133 and CYN1 showed typical characteristic peaks of the α-helical secondary structure with negative ellipticity at 222 and 208 nm (Figure 4A and 4B), suggested that their secondary structure was dominated by α-helix which was consistent with our design of four-helix bundle. Although the CD intensity decreased as the temperature increased from 25 °C to 95 °C, the negative peaks at 222 and 208 nm were retained, indicating that the helical structure of the protein sample at room temperature retained part of the secondary helical structure at high temperatures. Figure 4A and 4B further showed the CD spectrum of the initial 25 °C spectrum (black) compared with the spectrum measured at 95 °C (red) and at 25 °C after the melt (green). The spectra before and after refolding were very similar, indicating that the protein does refold once the temperature reduced, confirming that the folding was reversible. The CD-spectrum-derived helical contents according to the K2D3 web server is 52.2 % for D2-133 and 61.42 % for CYN1, suggesting that CYN1 has a more stable structural fold around the designed helical conformation (Figure 4E, F). However, this is still lower than 91% residues in helical regions for the designed protein.

**Figure 4.**
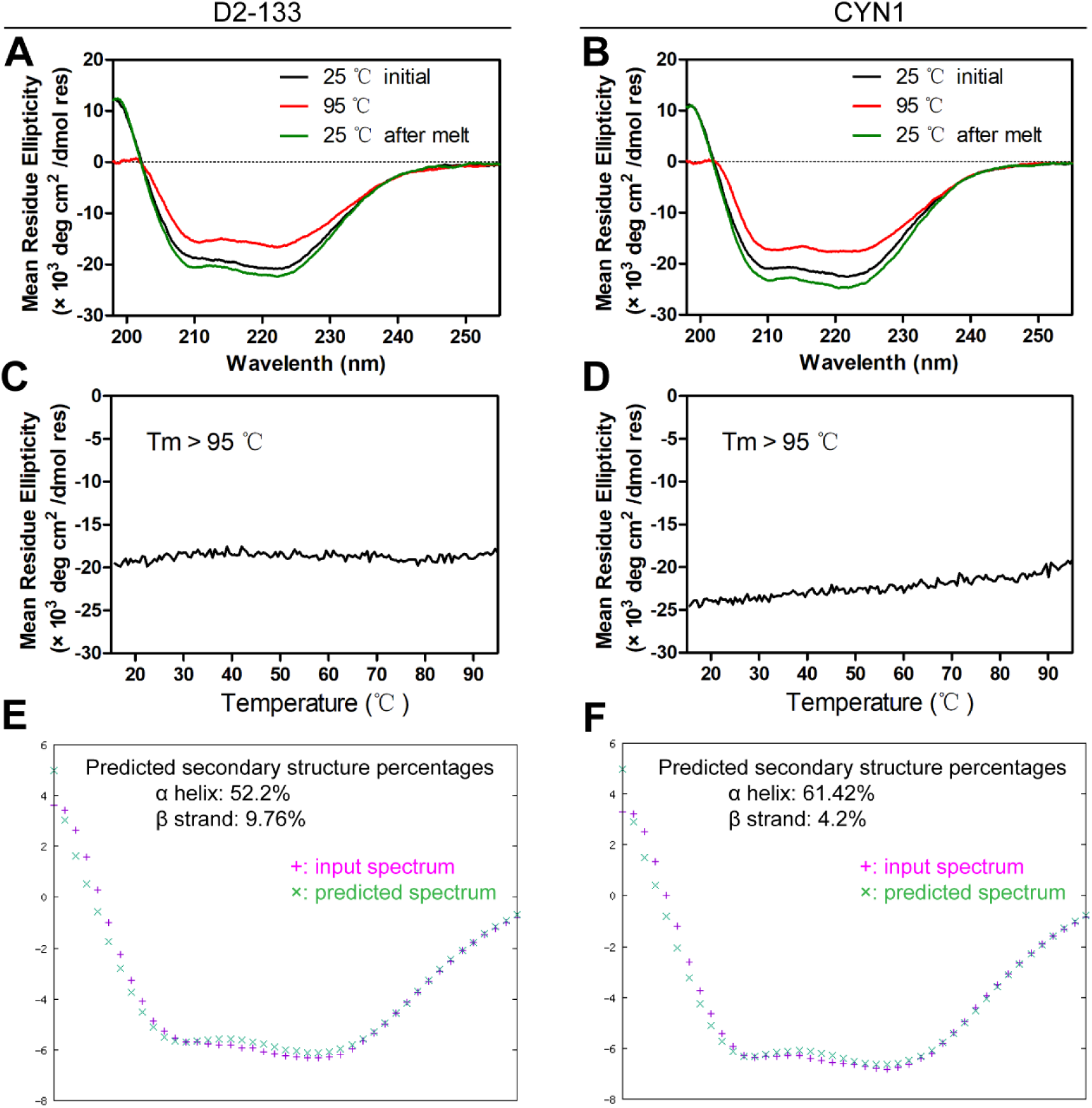
The Circular Dichroism Spectrum indicates high thermal and structural stability of D2-133 and CYN1. The Circular Dichroism Spectrum of D2-133 (**A**) and CYN1 (**B**) at 95 °C and 25 °C at native and refolded conditions as labeled. The CD signal of D2-133 (**C**) and CYN1 (**D**) at 222-nm wavelength, as a function of temperature. The predicted secondary structure contents of D2-133 (**E**) and CYN1(**F**) by K2D3 web server.

The CD signal strength as a function of temperature was monitored at 222 nm for D2-133 and CYN1 (Figure 4C and 4D). However, there is a lack of significant temperature-dependent changes suggesting that the melting temperatures of CYN1 and D2-133 were both greater than 95 °C. These results suggest that CYN1 exhibits the same high thermal stability as the designed H4 protein D2-133.

### The CYN1 showed broad-spectrum binding capability against RBD mutants from three different virus strains

To verify whether D2-133 and CYN1 bind broadly to RBDs of different SARS-CoV-2 strains, the RBDs of Omicron BA.1, XBB.1.16, and JN.1 virus strains were purchased for measuring binding affinity, and they had 13 mutations, 14 mutations, 27 mutations to the WT-RBD, respectively (Figure 5G). The results showed that CYN1 retains its activity in binding to RBDs from different virus strains (Figure 3E vs Figure 5A, 5B and 5C). Compared to the wild type RBD, the binding affinity of CYN1 to XBB.1.16 RBD was similar, at 17 nM, while the affinities of CYN1 to BA.1 and JN.1 RBDs were weakened, at 88 nM and 164 nM, respectively. The previously reported RBD binder AHB2 ^15^ was employed as a control, and its binding affinities to WT-RBD and BA.1 RBD were determined. As shown in Figure 5D, the binding affinity of AHB2 to WT-RBD was at 0.83 nM, consistent with the previous study ^15^. However, the affinity of AHB2 to the Omicron BA.1 RBD was only at 815 nM (Figure 5E), reduced by 1000-fold compared to the wildtype RBD (Figure 5F). By comparison, only a 4-fold reduction of binding of CYN1 to BA.1 relative to the binding to WT was observed (Figure 3E). It should be emphasized that the original design D2-133 does not display significant binding with the RBDs of BA.1, XBB.1.16 and JN.1 (Figure S5), despite of only 4 mutations away from CYN1.

**Figure 5.**
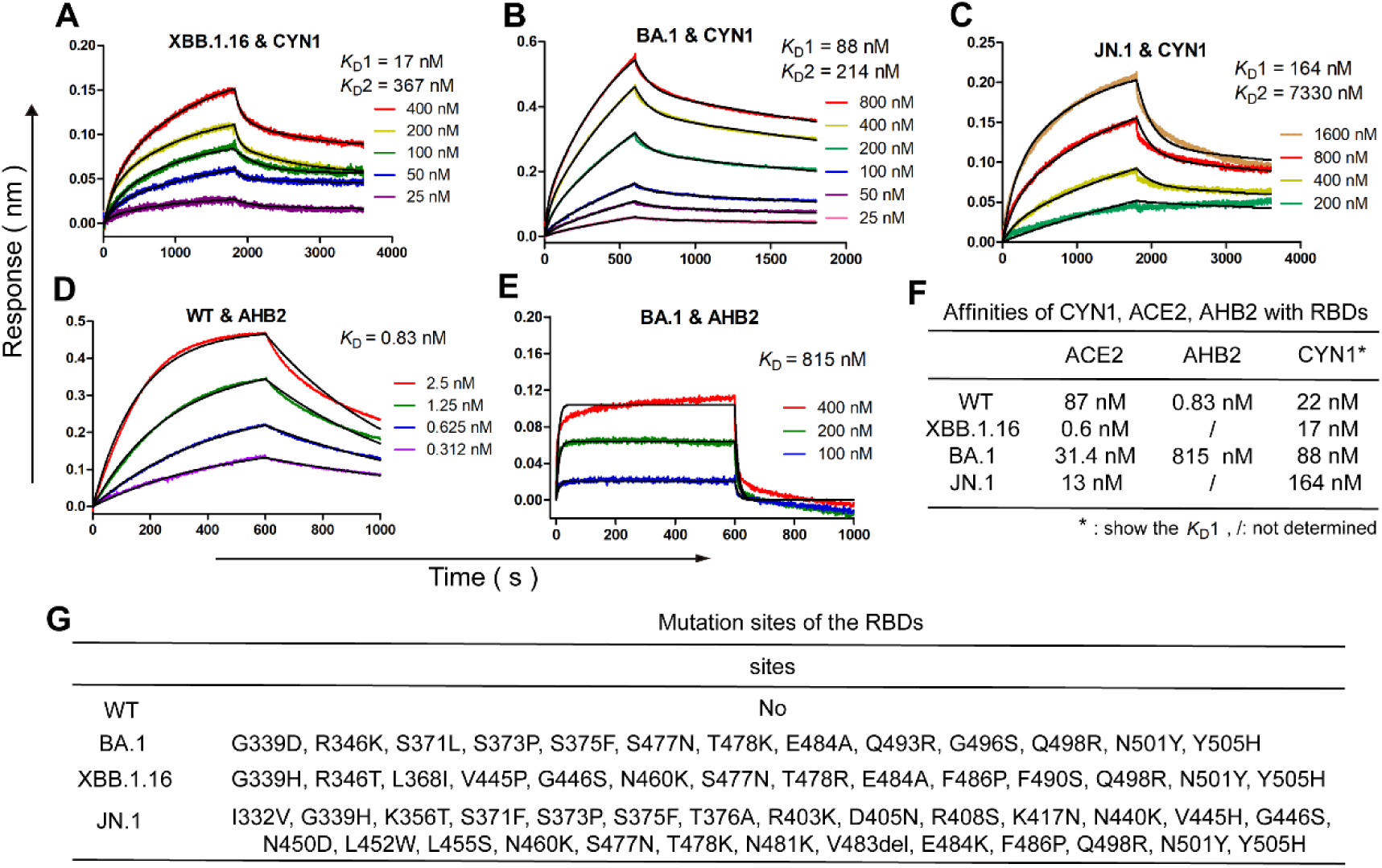
Broad spectrum measurement of binding to RBDs of other SARS-CoV 2 strains. Affinity of CYN1 with RBDs, fitted by 1:2 model to XBB.1.16 (**A**), BA.1 (**B**), JN.1 (**C**). Binding affinity of AHB2 fitted by 1:1 model to WT (**D**) and to BA.1 (**E**). (**F**) Table of Affinities of ACE2 ^7,47,50,51^, AHB2, and CYN1 with RBDs. (G) The mutation sites of the three Omicron RBDs to WT-RBD.

### Interpretation of binding free energy by molecular dynamics simulations and free energy calculations

To understand different behavior observed between D2-133 and CYN1, we performed molecular dynamics simulations of complexes of the WT-Spike RBD with ACE2, D2-133, E6, E9, and CYN1 and observed quite stable binding interfaces with the root mean square deviation (RMSD) values of the designed binding interfaces stable less than 3 Å for both 1-2 and 3-4 helices (Figures 6A and 6B). We then employed the ASGBIE method to calculate the interaction free energy between all residues within 10 Å of the binding interface and the other protein molecule. Fig. 6C shows a strong correlation with a Pearson correlation coefficient at 0.97 between experimental determined free energies and ASGBIE-calculated results according to simulated conformations. This consistency provides the basis for the following hot-spot analysis by computational alanine scanning.

**Figure 6.**
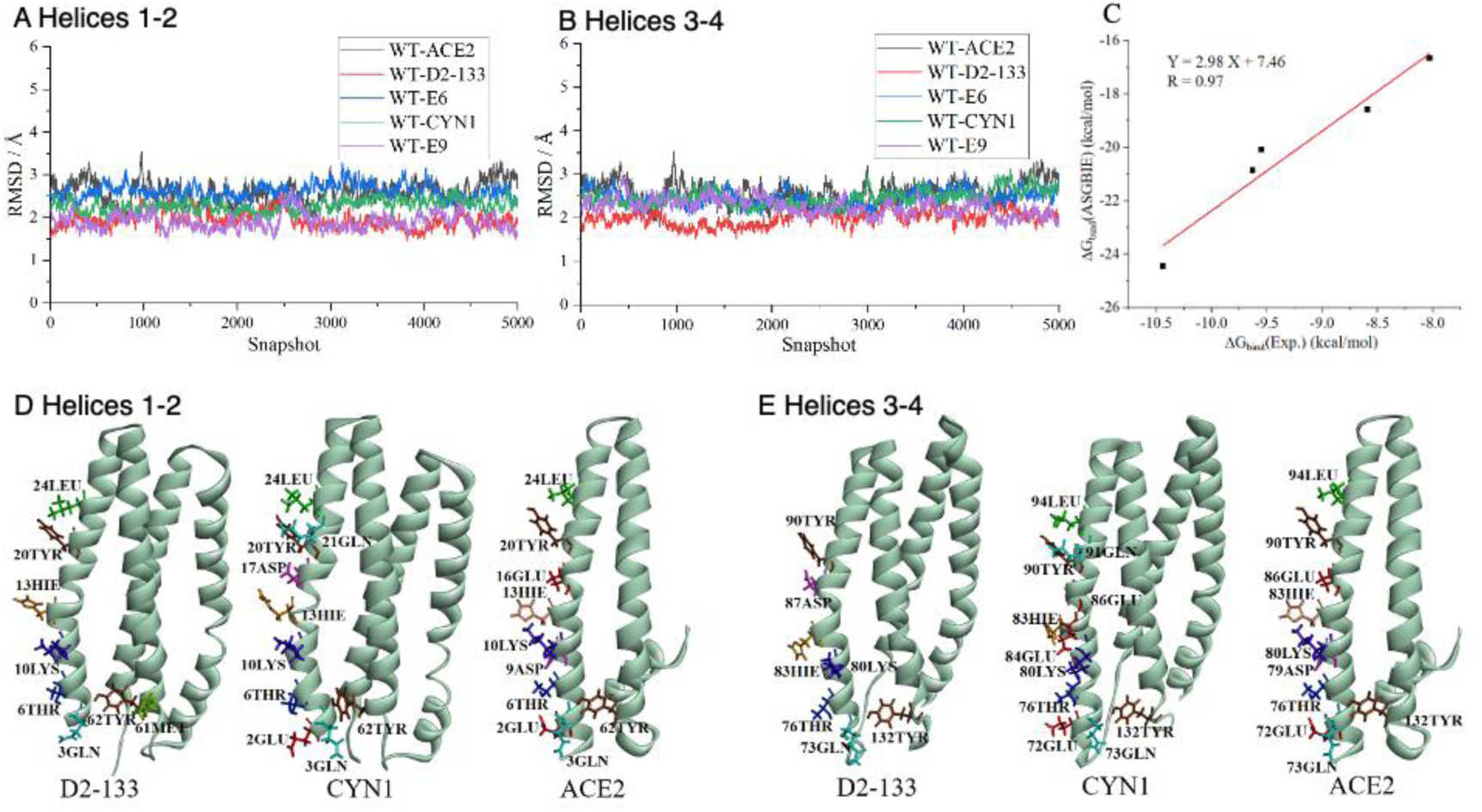
(A) Root-mean-squared deviation of helices 1-2 interface from the respective starting conformations by molecular dynamics simulations of ACE2, D2-133, E6, CYN1, and E9 in complex with a single WT-RBD of 5000 snap shots of 10 ns production runs after 40 ns equilibration run. (B) as in (A) but for helices 3-4 interface. (C) Calculated versus experimentally measured binding free energies of ACE2, D2-133, E6, CYN1, and E9 in complex with a single WT-RBD. (D) Comparison of hotspot residues identified by computational alanine scanning in helices 1-2 interface of D2-133 and CYN1 with the hotspot residues of ACE2 (E) As in (D) but for helices 3-4 interface. The sequence number in ACE2 was changed to match with those designed proteins for comparison.

### Hotspot residues and their implication on broad-spectrum capability

For binding of ACE2 to WT-RBD, we found a total of 8 hotspot residues, ranked by their contributions as follows: 20TYR, 13HIE, 10LYS, 16GLU, 62TYR, 3GLN, 2GLU, and 6THR. These hotspot residues were included as a part of fixed interfacial residues for designing ACE2 mimics. For our designed proteins, we have two surface regions mimicking the ACE2 interface to WT-RBD (helices 1-2 and helices 3-4). Thus, we performed two separated calculations to identify hotspots in two dual-helix regions. As shown in Figures 6D-E and Supplementary Tables S3-4), D2-133 has 6 out of 8 hotspot residues in the helices 1-2 interface and 6 out of 7 hotspot residues in helices 3-4 interface overlapping with ACE2 hotspot residues. By comparison, CYN1 has more hotspot residue interactions with WT-RBD with stronger binding affinities (10 for helix 1-2 and 11 for helix 3-4). Moreover, these hotspots overlap more with 8 ACE2 hotspot residues (7 for helix 1-2 and 8 for helix 3-4). This improved recovery of ACE2 hotspot residues by CYN1 explains its broader spectrum capability over D2-133.

## Discussion

SARS-CoV-2 has a fast mutation rate because it is an RNA virus with billions of hosts ^52,53^. Much research shows that the spike’s mutation was the most important factor for SARS-CoV-2 to escape the vaccine protection developed by pharmaceutical companies ^3,4,6,7,54^. The spike’s mutations also lead to resistance to antibody drugs because many antibody drugs were based on antigens on the spike proteins that are subsequently mutated ^55,56^. Here, we examined the possibility of designing a broad-spectrum protein by copying the same hACE2 interface that can bind to the RBDs of the different viral strains.

Despite the simplicity of the above idea, the design progress is not as straightforward as originally planned. In the first-round design, 2 out of 5 top-ranked designs did not have soluble expression, despite we have chosen the sequences with the lowest continuous hydrophobic patch area, indicating that the largest hydrophobic patch area is not necessarily the best indicator for solubility. Nevertheless, three of the five designs were expressed soluble and purified (Figure S1) and all bind with WT-RBD with micromolar binding affinity (Figure S2). Although the binding affinity is much lower than the binding between ACE2 and WT-RBD, the rate of success in achieving binding by OSCAR-Design is high (60%).

However, these three expressed proteins existed in aggregated forms on the chromatographic column (Figure S3). The difficulty of expression and aggregation faced here is likely due to the high hydrophobic patch area of the ACE2 binding interface and the need to maintain two large hydrophobic patches for our design with two ACE2 binding interfaces. To address this problem, we incorporated a disulfide bond in our design. All six designs are now expressed and soluble. Moreover, three of the six designs tested now exist in significant monomeric form with D2-133 purely monomeric, corresponding well to the fact that it has the smallest hydrophobic patch area (361Å^2^) and two with the largest hydrophobic patch areas (382 Å^2^ and 384 Å^2^) are all in multimeric form.

D2-133, however, continues to bind at the micromolar level. Moreover, it does not bind to RBDs of other SARS-CoV-2 strains. This indicates that the D2-133 interface grafted from ACE2 for RBD binding is only partially functional. That is, D2-133 must not have all the original ACE2 interfacial residues in the right place to interact with Spike RBD because it has a weak affinity to WT-RBD and no affinity to the RBDs of other viral strains.

D2-133 was improved by a factor of 13 in binding affinity to WT-RBD with one round of MACS selections. However, the only two mutations (K70E and K90R) in E6 are unexpectedly all outside the pre-defined ACE2-mimic interface region. The second round of MACS selections added two more mutations. This time both mutations are in the pre-defined ACE2 interfacial regions (L59T and L129T), but mutations are from hydrophobic leucine to more hydrophilic threonine. One would expect the opposite as less hydrophobic interface would lead to lower binding affinity.

To gain some understanding why these mutations yielded higher affinity and improved resistance against RBD mutations, we conducted molecular dynamics simulations on all systems to investigate the microstructural features of binding. Both the 1-2 and 3-4 helical regions were simulated as binding sites, respectively. The binding free energies of four-helix structures with WT-RBD obtained by using the ASGBIE method have a near perfect correlation to experimental measured results. A further hotspot analysis indicates that CYN1’s hotspot residues (both helices 1-2 and helices 2-4 regions) have more overlap with ACE2 hotspot residues than D2-133, suggesting the likely reason behind improved broad-spectrum capability than D2-133. CYN1 has not achieved a perfect broad-spectrum capability to constantly evolving SARS-CoV-2. This is because the binding affinity of CYN1 to other strains’ RBDs are lower than those of hACE2 ^7,57,58^. Moreover, there are many strains’ RBDs we have not tested yet due to the huge experiment cost. We had tried to determine the RBD-CYN1 complex structure, but the RBD-CYN1 complex was unstable at high concentration, which maybe contributed by the CYN1’s high hydrophobic area or the 1: 2 binding style to RBD. Nevertheless, we demonstrated that mimicking the small interface of ACE2 has the potential to achieve broad-spectrum capability. We expected that this can be improved with constantly improved AI-based protein design techniques so that the interface can be more faithfully captured.

## Author Contribution

**Jian Zhan and Yaoqi Zhou:** Conceptualization, Investigation, Supervise, Writing-review & editing. **Qinghui Nie:** Protein expression and purification, *E. coli* display to evolve ACE2 mimic’s affinity, Determined thermostability and affinity, Writing the original draft, Writing-review & editing. **Hongmei Yin**: Sequence design of the mimics, Writing the original draft. **Ke Chen**: Design pipeline development, backbone design of the ACE2 mimics, Writing the original draft. **Chao Zhao**: Performed computational simulations and binding analysis, Writing the original draft. **Weiwei Li:** sample preparation for structure determination. **Jianxun Qi**: Supervising, Writing review & editing**. Jinle Tang:** Supervise, Writing-review & editing. **John Z.H. Zhang:** Supervise, review & editing.

## Acknowledgements

The author would like to thank **Dr. Haitao Yang** and his doctoral student **Yuchi Lu** for their effort for helping determination of the RBD-CYN1 complex structure by Crystallization. This research paper was supported by the National Natural Science Foundation of China (Grant No. 92370202, 22333006, 92270001) and the National Key R&D Program of China (Grant No. 2021YFF1200400). We gratefully acknowledge the High Performance Computing Cluster at Shenzhen Bay Laboratory for participating in completing this study.

## Conflict of Interest

All authors declare no financial interest. Zhan and Zhou are the CEO and the chair of the scientific advisor board for Ribopeutic, respectively.

## Notes

### Competing Interest Statement

The authors have declared no competing interest.

### Summary of Updates

Authors' information update: correct the authors' order.

