## Supplementary Material for "Designing a broad-spectrum four-helix bundle targeting different strains of SARS-CoV-2 Spike Receptor Binding Domains with ACE2-like binding interface"

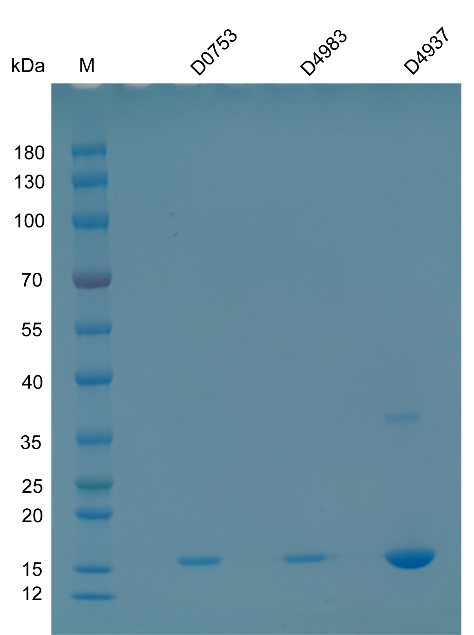

Figure S1. Three of the five first-round designed sequences (D0753, D4983, and D4937) were purified by Ni-NTA column (HisTrap FF crude 5 mL, Cytiva), then were separated on a 4-20% SDS-polyacrylamide gel and stained with Coomassie brilliant blue. M: protein ladder.

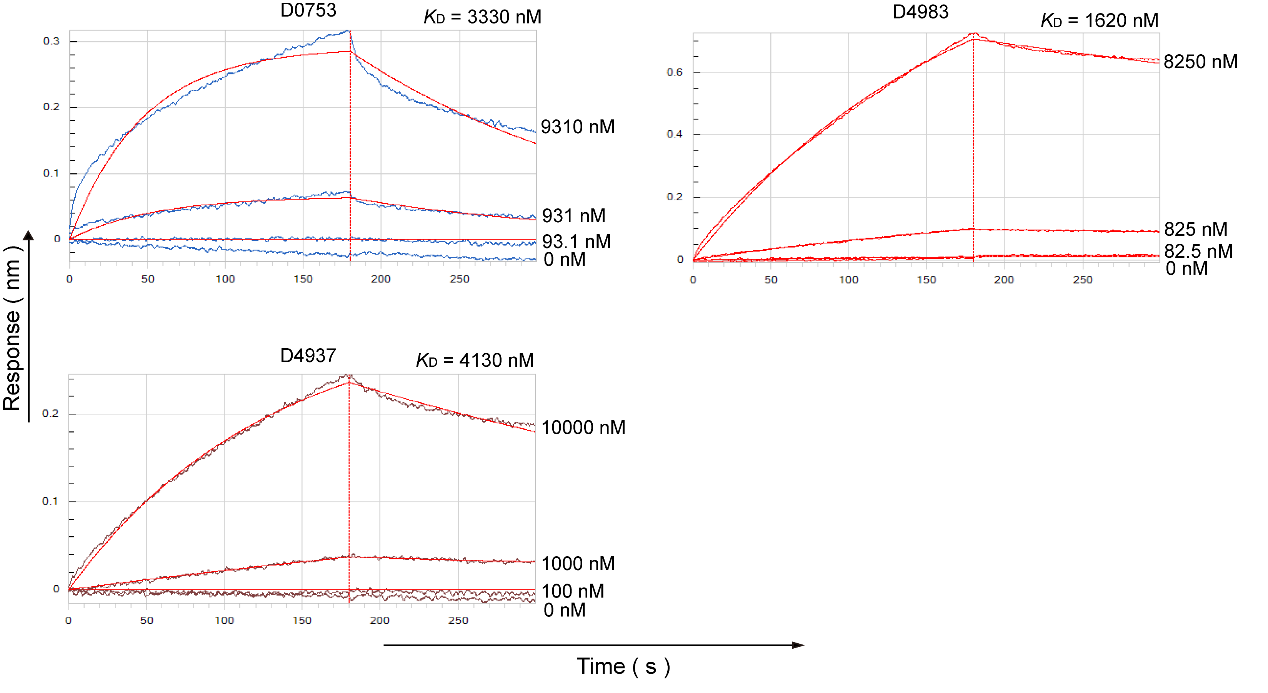

Figure S2. The binding affinities of three first-round designed proteins (D0753, D4983, and D4937) with the WT-RBD measured by biolayer interferometry.

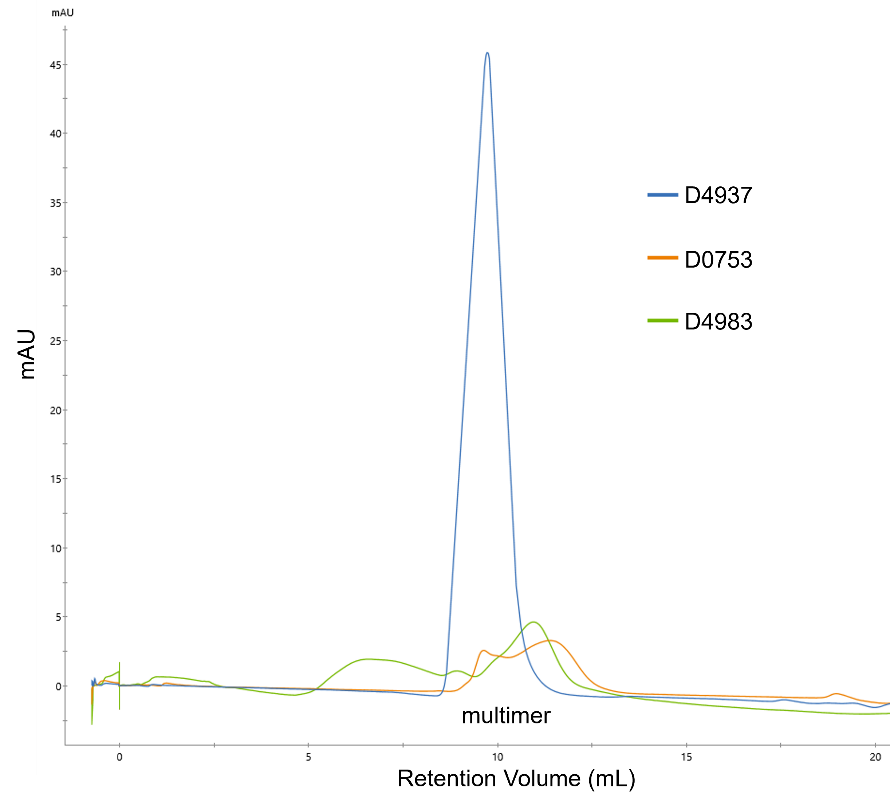

Figure S3. Size-exclusion chromatography (Superdex 75 Increase 10/300 GL, Cytiva) of three first-round designs (D0753, D4983, and D4937) on the ÄKTA pure™ 25M system (Cytiva).

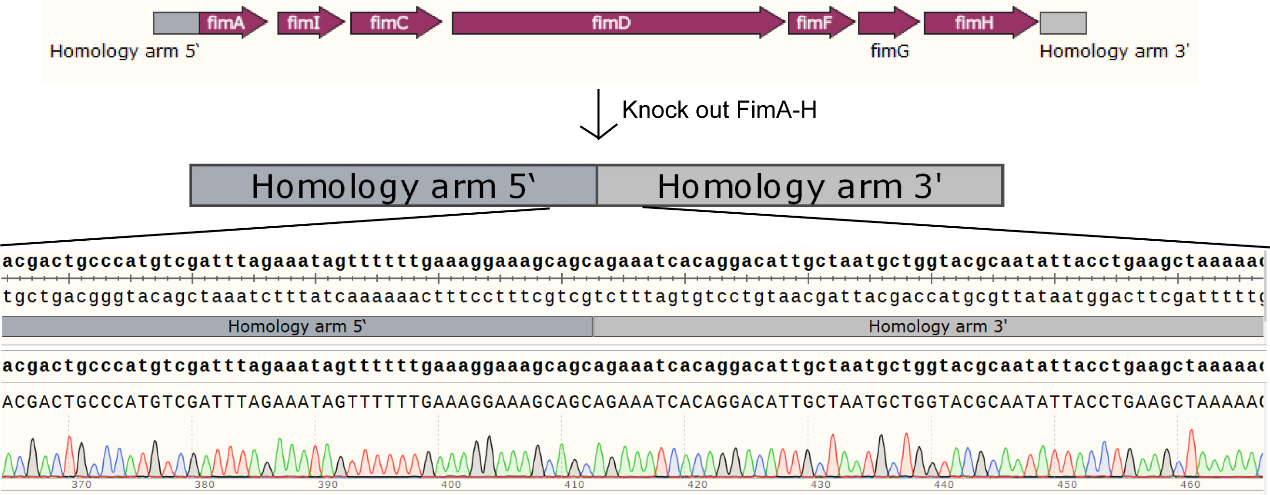

Figure S4. Sanger sequencing confirmed that *fim A-H* was knockout and *MG1655^ΔfimA-H^* strain was successfully constructed.

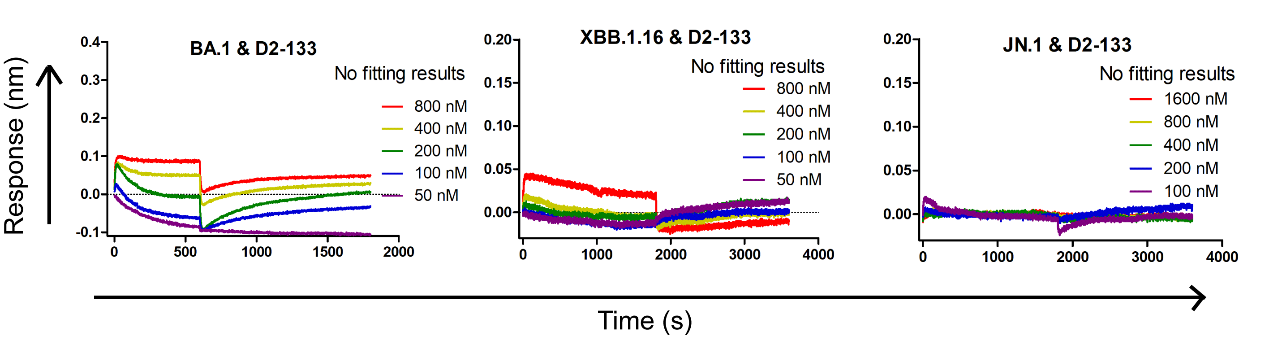

Figure S5. The lack of significant binding of D2-133 the RBDs of BA.1, XBB.1.16 and JN.1, as measured by the BLI.

**Table S1.** The protein sequences and the largest hydrophobic patch area in the first-round designs of H4 along with the experimental result (expressed or not expressed and measured binding affinity).

| ID | Sequences | $\boldsymbol{S}_{\boldsymbol{HPA}}$ (Å^2^) | Binding Affinity |
| --- | --- | --- | --- |
| D0064 | GEQLKTFADKFMHELEDIWYQAWLAVLDYQRNRTEEHRSNALNNLQKMFEFLDEQLRLLKMYGQYDKGNQQEQLNTFFDKFTHELEDIRYQAWLAYKNFKENMTDDHVNNAFNNLQKYREFLIEQLRLLLMYDFSQV | 375 | No Exp. |
| D0753 | GEQYNTFVDKFLHEFEDIWYQFQLAYQQYQENRTEEHRENFNNNLEKFYQFLMEQLRLFFMYGQHDKGTIQEQLNTFMDKFKHELEDILYQFQLARRNYQENKTDDSVKNAYNNAKKYAEFLLEQLRLFSMYDLSQV | 375 | 3330nM |
| D0420 | GEQFNTFLDKYLHEFEDIWYQAWLAYVQYQQNKTDSARSNTLNNFTKFWQFLDEQLRLFMMYGQYDQGTDQEQWETFMDKFKHELEDIRYQAWLAYKNFKENMTDDHVKNAFNNFQKYAEFLLEQFRLASMYDFNNI | 381 | No Exp. |
| D4937 | GEQLKTFADKFLHEAEDIYYQMVLAVQQYREQRTEENRKNVRNNILKFWQFLLEQLRLLEMYGQYDKGEIQEQLNTFFDKFKHELEDIKYQAWLAIKNFFENETDNHVDNALNNLQKYYNFLLEQLRLLLMYDFSKV | 373 | 4130nM |
| D4983 | GEQLKTFADKFLHEFEDIWYQTWLAVEDYRRNRTEEHRQNVVNNLQKFYQFLQEQLRLLKMYGQYDKGNVQEQFNTFLDKFTHELEDILYQFQLAYTQFVQNQTDNHVDNAWNNAEKYAQFLLEQLRLRLMYDFSQI | 382 | 1620nM |

**Table S2.** The protein sequences and the largest hydrophobic patch area in the second-round designs of H4 with a disulfide bond along with the experimental result on its structural form.

| Sequence ID | CYS Locations | $\boldsymbol{S}_{\boldsymbol{HPA}}$ | $\mathbf{Structure}$ |
| --- | --- | --- | --- |
| D2-133 | 1-133 | 361 | Monomer |
| D2-099 | 92-118 | 374 | partly monomeric |
| D2-128 | 22-48 | 379 | Multimeric |
| D2-477 | 25-44 | 380 | Mostly Monomer |
| D2-247 | 95-114 | 382 | Multimeric |
| D2-830 | 70-136 | 384 | Multimeric |

**Table S3.** The free energy (kcal/mol) of hot spots in helices 1-2 of D2-133, CYN1, and ACE2 when binding with RBD of WT SARS-CoV-2.

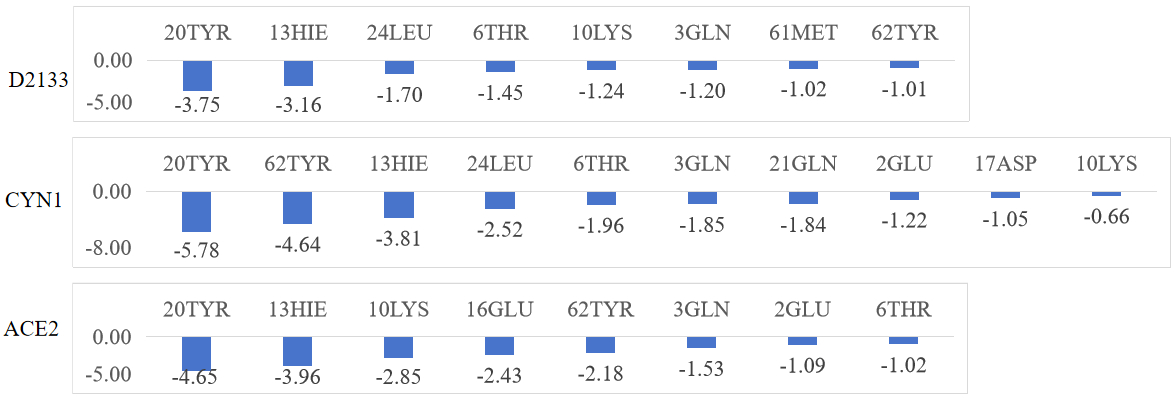

**Table S3.** The free energy (kcal/mol) of hot spots in RBD of WT SARS-CoV-2 when binding with helices 1-2 of D2-133, CYN1, and ACE2.

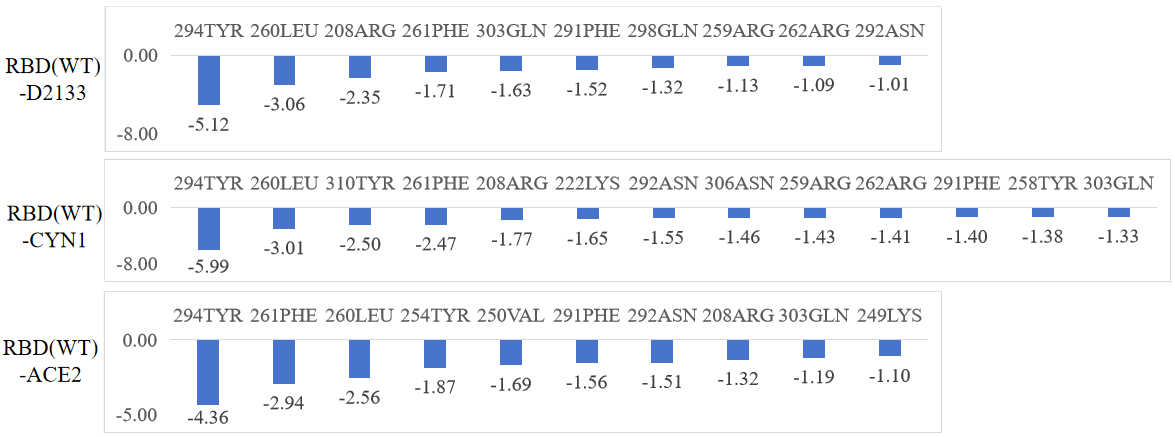

**Table S4.** The free energy (kcal/mol) of hot spots in helices 3-4 of D2-133, CYN1, and ACE2 when binding with RBD of WT SARS-CoV-2

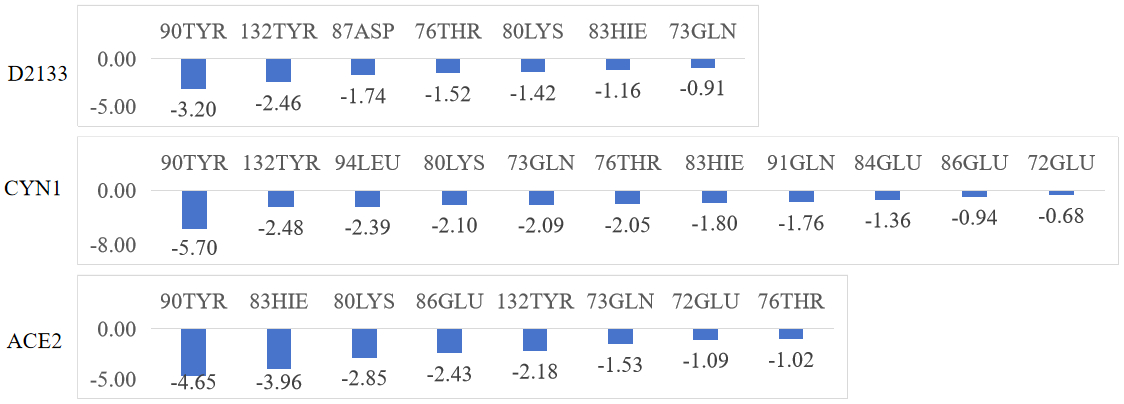

**Table S5.** The sequences of second-round designed proteins (D2), MACS-selected proteins in the first round (E-6 and E-9) and the second round (CYN1), and the control AHB2 protein.

| Name | Sequences |
| --- | --- |
| D2-133 | MCEQLETFADKFLHEAEDIWYQTWLAYVQYRQNKTSSQLQNFSNNLQKFKQFMDEQLRLLFMYGQMGKGKDQEQLDTFWDKFMHELEDIKYQFDLAYTQFQSNQTDSSASNAWNNAEKYYQFMLEQLRLLLMYCLNNV |
| D2-128 | MGEQLKTFADKYMHELEDILYQCLLAAENVRKNDTDENRRNFVNNLRKCWEFLKEQLRLLEMYGQYGKGRDQEQYNTFLDKFTHELEDTKYQTQLAIDQHESNITASSIDNALNNLLKYIDFMREQLRLRLMYDFNNI |
| D2-477 | MGEQLKTFADKYLHELEDIEYQARLCVQQYRENRTEEQRQNVANCLRKLWEFLMEQLRLLWMYGQYGKGNVQEQLNTFFDKFRHELEDIKYQAELALTQFYSNQTDDSVDNAWNNMLKYIEFMREQLRLLLMYDFNEV |
| D2-830 | MREQLVTFADKYMHELEDIYYQAWLAYVKYQQQNTSSNLQNFANNLTKLWQFLEEQLRLLWMYGQYGKGKCQEQLNTFFDKFRHELEDIRYQAWLAIKNFHENKTDDSSKNAWNNMQKFIDFMREQLRLLLMYDFSCV |
| D2-099 | MREQLVTFTDKYLHELEDIWYQAWLAVEDYRRDRKEDNRKNVKNNLQKLLEFLDEQRRLLEMYGQYGKGNVQEQFTTFEDKRQHELEDTLYQCRLAYTQFVSNMTDSSVDNAWNNMEKCIQFLREQLRLRLMYDFNEI |
| D2-247 | MGEQLKTFADKFLHEAEDIWYQTWLAVEQHRQNDTDDTRKNVLNNLQKFYEFMMEQLRLLWMYGQYGKGRDQEQLTTFFDKFTHELEDILYQFQLCYTQFKSNMTQDSVDNAWNCAEKFYQFMLEQLRLLLMYDFNNV |
| E-6 | MCEQLETFADKFLHEAEDIWYQTWLAYVQYRQNKTSSQLQNFSNNLQKFKQFMDEQLRLLFMYGQMGKGEDQEQLDTFWDKFMHELEDIRYQFDLAYTQFQSNQTDSSASNAWNNAEKYYQFMLEQLRLLLMYCLNNV |
| E-9 | MCEQLETFADKFLHEAEDIWYQTWLAYVQYRQNKTSSQLQNFSNNLQKFKQFMDEQLRLLFMYGQMGKGKDQEQLDTFWDKFMHELEDIKYQFDLAYTQFQSIQTDSSASNAWNNAEKYYQFMLGQLRLLLMYCLNNV |
| CYN1 | MCEQLETFADKFLHEAEDIWYQTWLAYVQYRQNKTSSQLQNFSNNLQKFKQFMDEQLRTLFMYGQMGKGEDQEQLDTFWDKFMHELEDIRYQFDLAYTQFQSNQTDSSASNAWNNAEKYYQFMLEQLRTLLMYCLNNV |
| AHB2 | MELEEQVMHVLDQVSELAHELLHKLTGEELERAAYFNWWATEMMLELIKSDDEREIREIEEEARRILEHLEELARK |

**Table S6.** DNA sequences of first-round (D-) and second-round designed proteins (D2-), MACS-selected proteins in the first round (E-6 and E-9) and the second round (CYN1) as well as sequences for primers and donor DNAs.

| Name | DNA sequences (5’-3’) |
| --- | --- |
| D0064 | ATGGGCGAACAGCTGAAAACCTTTGCGGATAAATTTATGCATGAGCTGGAAGACATTTGGTACCAGGCGTGGCTGGCGGTGCTGGATTATCAGCGCAATCGCACCGAAGAACATCGCAGCAATGCGCTGAATAATCTGCAGAAAATGTTTGAGTTTCTGGACGAACAGCTGCGCCTGCTGAAAATGTATGGCCAGTATGATAAAGGCAATCAGCAGGAACAGCTGAATACCTTCTTTGATAAATTCACCCACGAGCTGGAAGATATTCGCTATCAGGCGTGGCTGGCGTATAAAAATTTTAAAGAAAATATGACCGACGACCACGTGAACAATGCGTTTAATAATCTGCAGAAATACCGCGAATTTCTGATCGAACAGCTGCGCCTGCTGCTGATGTATGATTTTAGCCAGGTG |
| D0753 | ATGGGCGAACAGTATAATACCTTTGTGGATAAATTTCTGCATGAGTTCGAGGATATCTGGTATCAGTTTCAGCTGGCGTATCAGCAGTATCAGGAAAATCGCACCGAAGAACATCGCGAAAATTTTAATAATAACCTGGAGAAGTTCTACCAGTTCCTGATGGAACAGCTGCGCCTGTTCTTTATGTATGGCCAGCATGATAAAGGCACCATTCAGGAACAGCTGAATACCTTTATGGATAAATTTAAACACGAGCTGGAAGACATCCTGTATCAGTTTCAGCTGGCCCGCCGCAATTATCAGGAAAATAAAACCGATGATAGCGTGAAAAATGCGTATAATAATGCGAAAAAGTACGCGGAGTTTCTGCTGGAACAGCTGCGCCTGTTTAGCATGTATGATCTGAGCCAGGTG |
| D0420 | ATGGGCGAACAGTTTAATACCTTTCTGGATAAATATCTGCACGAGTTCGAGGATATTTGGTATCAGGCGTGGCTGGCGTATGTGCAGTATCAGCAGAATAAAACCGATAGCGCCCGTAGCAATACCCTGAATAATTTTACCAAATTTTGGCAGTTCCTGGACGAACAGCTGCGCCTGTTTATGATGTATGGCCAGTATGATCAGGGCACCGATCAGGAACAGTGGGAAACCTTTATGGATAAATTTAAGCACGAGCTGGAGGATATCCGCTATCAGGCGTGGCTGGCGTATAAAAATTTTAAAGAAAATATGACCGACGACCACGTGAAGAATGCGTTTAATAATTTTCAGAAATACGCGGAGTTCCTGCTGGAACAGTTTCGCCTGGCGAGCATGTATGATTTTAATAATATT |
| D4937 | ATGGGCGAACAGCTGAAAACCTTTGCGGATAAATTTCTGCATGAAGCGGAAGATATTTACTACCAGATGGTGCTGGCGGTGCAGCAGTATCGCGAACAGCGCACCGAAGAAAATCGCAAAAATGTGCGCAATAATATCCTGAAGTTTTGGCAGTTTCTGCTGGAACAGCTGCGCCTGCTGGAAATGTATGGCCAGTATGATAAAGGCGAAATTCAGGAACAGCTGAATACCTTCTTTGACAAATTTAAGCACGAACTGGAGGACATCAAGTACCAGGCGTGGCTGGCGATTAAAAATTTCTTTGAAAATGAGACTGACAACCACGTGGACAACGCGCTGAATAATCTGCAGAAATATTATAACTTCCTGCTGGAGCAGCTGCGCCTGCTGCTGATGTATGATTTTAGCAAAGTG |
| D4983 | ATGGGCGAACAGCTGAAAACCTTTGCGGATAAATTTCTGCATGAATTCGAAGACATTTGGTACCAGACCTGGCTGGCGGTGGAAGATTATCGCCGCAATCGCACCGAAGAACATCGCCAGAATGTGGTGAATAATCTGCAGAAATTTTACCAGTTTCTGCAGGAGCAGCTGCGCCTGCTGAAAATGTATGGCCAGTATGATAAAGGCAATGTGCAGGAACAGTTTAATACCTTTCTGGATAAATTCACCCACGAGCTGGAAGATATTCTGTATCAGTTTCAGCTGGCGTATACCCAGTTTGTGCAGAATCAGACCGATAATCATGTGGATAATGCGTGGAATAATGCGGAAAAATATGCGCAGTTTCTGCTGGAACAGCTGCGCCTGCGCCTGATGTATGATTTTAGCCAGATT |
| D2-133 | ATGTGTGAACAGCTGGAAACCTTTGCAGATAAATTTCTGCATGAGGCCGAAGATATTTGGTATCAGACCTGGCTGGCCTATGTTCAGTATCGTCAGAATAAAACCAGCAGCCAGCTGCAGAATTTTAGCAATAATCTGCAGAAATTCAAGCAGTTCATGGACGAGCAGCTGCGCCTGCTGTTTATGTATGGTCAGATGGGTAAAGGTAAAGACCAGGAACAGCTGGATACCTTTTGGGATAAATTTATGCACGAGCTGGAGGATATCAAGTATCAGTTTGACCTGGCCTATACCCAGTTTCAGAGCAATCAGACCGATAGCAGCGCAAGCAATGCATGGAATAATGCAGAAAAATACTACCAGTTCATGCTGGAACAGCTGCGTCTGCTGCTGATGTATTGTCTGAATAATGTT |
| D2-128 | ATGGGTGAACAGCTGAAAACCTTTGCAGATAAATATATGCACGAGCTGGAGGACATTCTGTATCAGTGTCTGCTGGCAGCAGAAAATGTTCGTAAAAATGATACCGACGAGAACCGCCGTAATTTCGTGAATAATCTGCGCAAGTGTTGGGAGTTCCTGAAAGAACAGCTGCGCCTGCTGGAAATGTATGGTCAGTATGGTAAAGGTCGTGATCAGGAACAGTATAATACCTTTCTGGATAAGTTCACCCATGAGCTGGAGGATACCAAATATCAGACCCAGCTGGCAATTGATCAGCATGAAAGCAATATTACCGCCAGCAGCATTGATAATGCACTGAATAATCTGCTGAAGTACATCGACTTCATGCGTGAACAGCTGCGCCTGCGTCTGATGTATGATTTTAATAATATC |
| D2-477 | ATGGGTGAACAGCTGAAAACCTTTGCAGATAAATATCTGCACGAGCTGGAAGACATTGAGTATCAGGCACGTCTGTGTGTTCAGCAGTATCGTGAAAATCGTACCGAAGAACAGCGTCAGAATGTTGCAAATTGTCTGCGTAAACTGTGGGAATTTCTGATGGAACAGCTGCGTCTGCTGTGGATGTATGGTCAGTATGGTAAAGGTAATGTGCAGGAGCAGCTGAATACCTTTTTTGACAAGTTTCGCCATGAGCTGGAAGACATCAAATACCAGGCAGAACTGGCACTGACCCAGTTTTATAGCAATCAGACCGATGATAGCGTTGACAATGCATGGAATAATATGCTGAAGTATATTGAGTTCATGCGCGAACAGCTGCGCCTGCTGTTAATGTATGATTTTAATGAAGTG |
| D2-830 | ATGCGTGAACAGCTGGTTACCTTTACCGATAAATATCTGCATGAGCTGGAAGACATTTGGTACCAGGCATGGCTGGCAGTTGAAGATTATCGTCGTGATCGTAAAGAAGACAATCGCAAAAATGTGAAAAACAACCTGCAGAAGCTGCTGGAATTTCTGGATGAACAGCGTCGTCTGCTGGAAATGTATGGTCAGTATGGTAAAGGTAATGTGCAGGAGCAGTTTACCACCTTTGAAGACAAACGTCAGCATGAGCTGGAAGACACCCTGTATCAGTGTCGTCTGGCATATACCCAGTTTGTTAGCAATATGACCGATAGCAGCGTTGATAATGCATGGAATAATATGGAGAAGTGCATCCAGTTCCTGCGCGAGCAGCTGCGTTTACGTTTAATGTATGATTTTAACGAGATC |
| D2-099 | ATGGGTGAACAGCTGAAAACCTTTGCAGATAAATTTCTGCACGAGGCCGAGGATATTTGGTATCAGACCTGGCTGGCAGTTGAACAGCATCGTCAGAATGATACCGATGATACCCGTAAAAATGTGCTGAATAATCTGCAGAAATTCTACGAGTTCATGATGGAGCAGCTGCGTCTGCTGTGGATGTATGGTCAGTATGGTAAAGGTCGTGATCAGGAACAGCTGACCACCTTTTTTGATAAATTTACCCACGAGCTGGAGGACATCCTGTATCAGTTTCAGCTGTGTTACACCCAGTTTAAGAGCAATATGACCCAGGATAGCGTTGATAATGCATGGAATTGTGCAGAAAAATTCTACCAGTTCATGCTGGAGCAGCTGCGCCTGCTGTTAATGTATGATTTTAATAATGTG |
| D2-247 | ATGCGTGAACAGCTGGTTACCTTTGCAGATAAATATATGCACGAGCTGGAAGACATTTACTACCAGGCATGGCTGGCATACGTTAAATATCAGCAGCAGAATACCAGCAGCAATCTGCAGAATTTTGCAAATAATCTGACCAAGCTGTGGCAGTTTCTGGAAGAACAGCTGCGTCTGCTGTGGATGTATGGTCAGTATGGTAAAGGTAAATGCCAGGAGCAGCTGAATACCTTTTTTGACAAATTCCGCCATGAGCTGGAAGACATTCGTTACCAGGCATGGCTGGCAATTAAAAATTTTCACGAGAACAAGACCGACGACAGCAGCAAAAATGCCTGGAATAATATGCAGAAATTCATCGACTTCATGCGCGAACAGCTGCGTCTGCTGCTGATGTATGATTTTAGCTGTGTT |
| E-6 | ATGTGTGAACAGCTGGAAACCTTTGCAGATAAATTTCTGCATGAGGCCGAAGATATTTGGTATCAGACCTGGCTGGCCTATGTTCAGTATCGTCAGAATAAAACCAGCAGCCAGCTGCAGAATTTTAGCAATAATCTGCAGAAATTCAAGCAGTTCATGGACGAGCAGCTGCGCCTGCTGTTTATGTATGGTCAGATGGGTAAAGGTGAAGACCAGGAACAGCTGGATACCTTTTGGGATAAATTTATGCACGAGCTGGAGGATATCAGGTATCAGTTTGACCTGGCCTATACCCAGTTTCAGAGCAATCAGACCGATAGCAGCGCAAGCAATGCATGGAATAATGCAGAAAAATACTACCAGTTCATGCTGGAACAGCTGCGTCTGCTGCTGATGTATTGTCTGAATAATGTT |
| E-9 | ATGTGTGAACAGCTGGAAACCTTTGCAGATAAATTTCTGCATGAGGCCGAAGATATTTGGTATCAGACCTGGCTGGCCTATGTTCAGTATCGTCAGAATAAAACCAGCAGCCAGCTGCAGAATTTTAGCAATAATCTGCAGAAATTCAAGCAGTTCATGGACGAGCAGCTGCGCCTGCTGTTTATGTATGGTCAGATGGGTAAAGGTAAAGACCAGGAACAGCTGGATACCTTTTGGGATAAATTTATGCACGAGCTGGAGGATATCAAGTATCAGTTTGACCTGGCCTATACCCAGTTTCAGAGCATTCAGACCGATAGCAGCGCAAGCAATGCATGGAATAATGCAGAAAAATACTACCAGTTCATGCTGGGACAGCTGCGTCTGCTGCTGATGTATTGTCTGAATAATGTT |
| CYN1 | ATGTGTGAACAGCTGGAAACCTTTGCAGATAAATTTCTGCATGAGGCCGAAGATATTTGGTATCAGACCTGGCTGGCCTATGTTCAGTATCGTCAGAATAAAACCAGCAGCCAGCTGCAGAATTTTAGCAATAATCTGCAGAAATTCAAGCAGTTCATGGACGAGCAGCTGCGCACGCTGTTTATGTATGGTCAGATGGGTAAAGGTGAAGACCAGGAACAGCTGGATACCTTTTGGGATAAATTTATGCACGAGCTGGAGGATATCAGGTATCAGTTTGACCTGGCCTATACCCAGTTTCAGAGCAATCAGACCGATAGCAGCGCAAGCAATGCATGGAATAATGCAGAAAAATACTACCAGTTCATGCTGGAACAGCTGCGTACGCTGCTGATGTATTGTCTGAATAATGTT |
| AHB2  (binding positive control) | ATGGAACTGGAGGAACAGGTAATGCACGTTCTGGACCAGGTGTCTGAACTGGCCCACGAGCTGCTGCACAAACTGACCGGTGAAGAACTGGAACGTGCGGCTTACTTCAACTGGTGGGCTACGGAGATGATGCTGGAGCTGATCAAATCTGACGACGAACGCGAAATCCGTGAAATCGAGGAAGAAGCGCGTCGTATCCTGGAGCACCTGGAAGAACTGGCGCGCAAA |
| P133SF  (Primer of subclone D2-133 variants into pET-21 a (+)) | ACTTTAAGAAGGAGATATACATATGTGTGAACAGCTGGAAAC |
| P133SR  (Primer of subclone D2-133 variants into pET-21 a (+)) | TGGTGGTGGTGGTGCTCGAGCTCGAGAACATTATTCAGACAATAC |
| PD133F (Error prone PCR primer for D2-133) | gtgcgactagAGGTACCATGTGTGAACAGCTGGAAAC |
| PD133R  (Error prone PCR primer for D2-133) | gaaataagcttttgttcggatccGTGGTGGTGGTGGTGG |
| donorF  (PCR primer for donor DNA) | caaaaagcatctaactgtttgat |
| donor (PCR primer for donor DNA) | gttcaaaggttactttgacctg |
| FimtsF (Primer for check the donor DNA in the genome) | ggcggagcgtggtgcagatactcg |
| FimtsR (Primer for check the donor DNA in the genome) | gaccgccgccgggattatcagtgc |
| Homology arm fragments  (The donor DNA for knock out fimbriae of MG1655) | caaaaagcatctaactgtttgatatgtaaattatttctattgtaaattaatttcacatcacctccgctatatgtaaagctaacgtttctgtggctcgacgcatcttcctcattcttctctccaaaaaccacctcatgcaatataaacatctataaataaagataacaatagaatattaagccaacaaataaactgaaaaagtttgtccgcgatgctttcctctatgagtcaaaatggccccaaatgtttcatcttttgggggaaaactgtgcagtgttggcagtcaaactcgtttacaaaacaaagtgtacagaacgactgcccatgtcgatttagaaatagttttttgaaaggaaagcagcagaaatcacaggacattgctaatgctggtacgcaatattacctgaagctaaaaacctgcacgttagccctttgtaggccagataagacgcgtcagcgtcgcatctggcataaacaaagcgcactttgctggtctgttcccctcaccctaaccctctccccggaggggcgaggggactgtccgggcacatttttagactttgtcatcagtctgagcctgccattggcaggctctggtgtccttttacgctaccatgctaataatcagcacaataatcagcccaaccacggagttgaccagctccagcagaccccaggttttcaacgtgtcttttactgacaggtcaaagtaacctttgaac |
